# Antibiotic susceptibility of bacterial colonies: An assay and experiments with *Staphylococcus aureus*

**DOI:** 10.1101/075515

**Authors:** Xinxian Shao, Justin Kim, Ha Jun Jeong, Bruce Levin

## Abstract

A method is presented to evaluate *in vitro* the efficacy of antibiotics to treat bacteria growing as discrete colonies on surfaces and the contribution of the colony structure to the antibiotic susceptibility of bacteria. Using this method, we explored the relative efficacy of six bactericidal and three bacteriostatic antibiotics to inhibit the growth and kill *Staphylococcus aureus* colonies of different sizes, densities and ages. As measured by the reduction in viable cell density relative to untreated controls, of the bactericidal drugs tested ciprofloxacin and gentamicin were most effective. By this criteria, ampicillin was more effective than oxacillin. Daptomycin and vancomycin were virtually ineffective for treating *S. aureus* growing as colonies. The bacteriostatic antibiotic tested, tetracycline, linezolid and erythromycin were all able to prevent the growth of *S. aureus* colonies and did so even more effectively than daptomycin, which is highly bactericidal in liquid culture. The results of these experiments and other observations suggest that relative inefficacy of oxacillin, vancomycin and daptomycin to kill *S. aureus* in colonies is due to the density and physiological state of the bacteria rather than the inability of these drugs to penetrate the colonies. The methods developed here are general and can be used to explore the efficacy of antibiotics to treat bacteria growing in biofilms as well as discrete colonies.

## Introduction

The rational (as opposed to purely empirical) approach to the design of antibiotic treatment regimes is based on estimates of the changes in the serum concentration of the drugs following their administration, pharmacokinetics (PK), and the relationship between the concentration of the drug and the rates of growth and death of the target bacteria, pharmacodynamics (PD) (1). Almost all we know about the PDs of antibiotics and bacteria is from *in vitro* studies of planktonic cells maintained in well-agitated liquid cultures (2-5) and the theoretical analog of these culture conditions, mass-action mathematical models (6–9). Under these conditions all the bacteria in a population have equal access to each other as well as resources, wastes, and allopathic agents like antibiotics.

In the real world of infections, bacteria are more likely to live in physically structured habitats, embedded in polysaccharide matrices known as biofilms (10–12) or as discrete colonies on the surfaces of tissues or within semi-solids (13–15). Under these conditions, the individual cells of bacterial population vary in their access to each other, nutrients, wastes and antibiotics. How does this reality of physically structured habitats affect the pharmacodynamics of antibiotics and thereby the rational design of antibiotic treatment protocols?

This question has been addressed and answers have been obtained for biofilms (11, 16–22). In this report, we consider the pharmacodynamics of antibiotics and bacteria growing as discrete colonies. We present a method to quantitatively evaluate the antibiotic susceptibility of bacteria growing as colonies on surfaces and compare their susceptibility to planktonic bacteria of the same densities and physiological state. Using this method, we explore the susceptibility of *Staphylococcus aureus* Newman colonies of different ages and sizes to six bactericidal and three bacteriostatic antibiotics. The results of our study indicate substantial variation in the efficacy of the tested antibiotics to treat *S. aureus* maintained as colonies. Antibiotics that are effective in killing *S. aureus* in liquid culture are virtually ineffective when these bacteria are growing as colonies.

## Material and Methods

### Media

The Mueller-Hinton II (MHII) medium used in liquid and agar cultures was Cation Adjusted MHII Becton Dickinson, (Franklin Lakes, NJ, USA).

Lysogeny Broth (LB) Becton Dickinson, (Franklin Lakes, NJ, USA) was used to prepare LB agar plates for sampling.

### Bacteria

*Staphylococcus aureus* Newman (generously provided by William Shafer) was used in this investigation. This strain has a clinical origin, remains virulent and has been extensively used in studies of staphylococcal pathogenicity (23).

### MIC Estimation

MICs were estimated in vitro using the serial dilution procedure similar to that described by (24) with modifications; we obtained these estimates in a standard MHII broth with inoculates of 5×10^7^cells/ml as well a the standard 5×10^5^cells/ml. In our serial dilution plates, we also used different starting concentrations of the antibiotics to obtain more precise estimates of the MICs. The results are shown in Table 1.

**Table 1.**
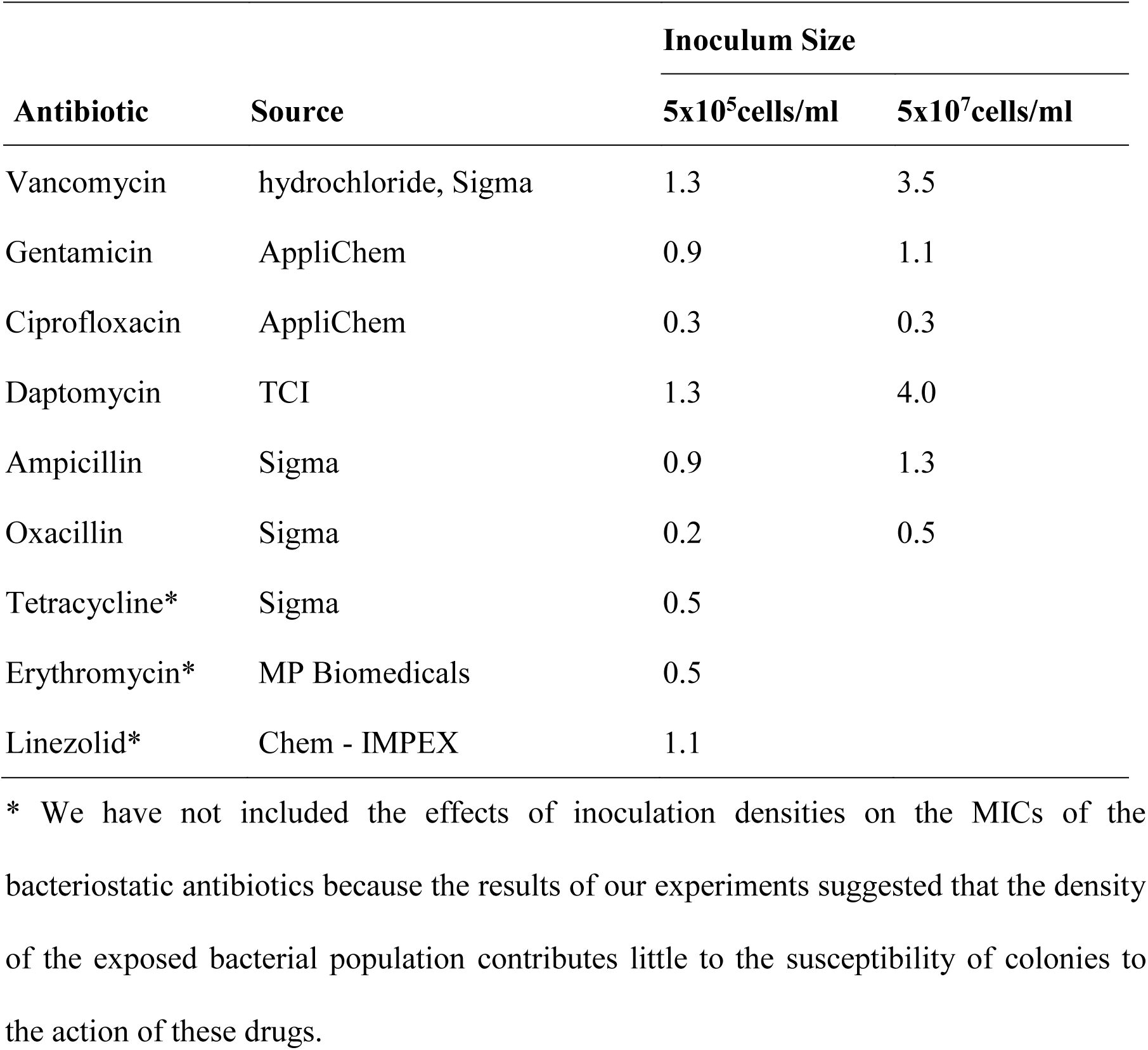
The MICs of 9 antibiotics in MHII broth of different concentrations and cultures initiated with different numbers of viable cells, unit of the concentrations is in µg/ml.

| Antibiotic | Source | Inoculum Size |  |
| --- | --- | --- | --- |
| | | $5 \times 10^5 \text{ cells/ml}$ | $5 \times 10^7 \text{ cells/ml}$ |
| Vancomycin | hydrochloride, Sigma | 1.3 | 3.5 |
| Gentamicin | AppliChem | 0.9 | 1.1 |
| Ciprofloxacin | AppliChem | 0.3 | 0.3 |
| Daptomycin | TCI | 1.3 | 4.0 |
| Ampicillin | Sigma | 0.9 | 1.3 |
| Oxacillin | Sigma | 0.2 | 0.5 |
| Tetracycline* | Sigma | 0.5 |  |
| Erythromycin* | MP Biomedicals | 0.5 |  |
| Linezolid* | Chem - IMPEX | 1.1 |  |

\* We have not included the effects of inoculation densities on the MICs of the bacteriostatic antibiotics because the results of our experiments suggested that the density of the exposed bacterial population contributes little to the susceptibility of colonies to the action of these drugs.

### Procedure for the Colony Assay for Antibiotic Efficacy and Liquid Culture Controls

A diagram of the method used to prepare the colonies, expose the bacteria to antibiotics and estimate viable cell densities is presented in Fig. 1.

**Figure 1.**
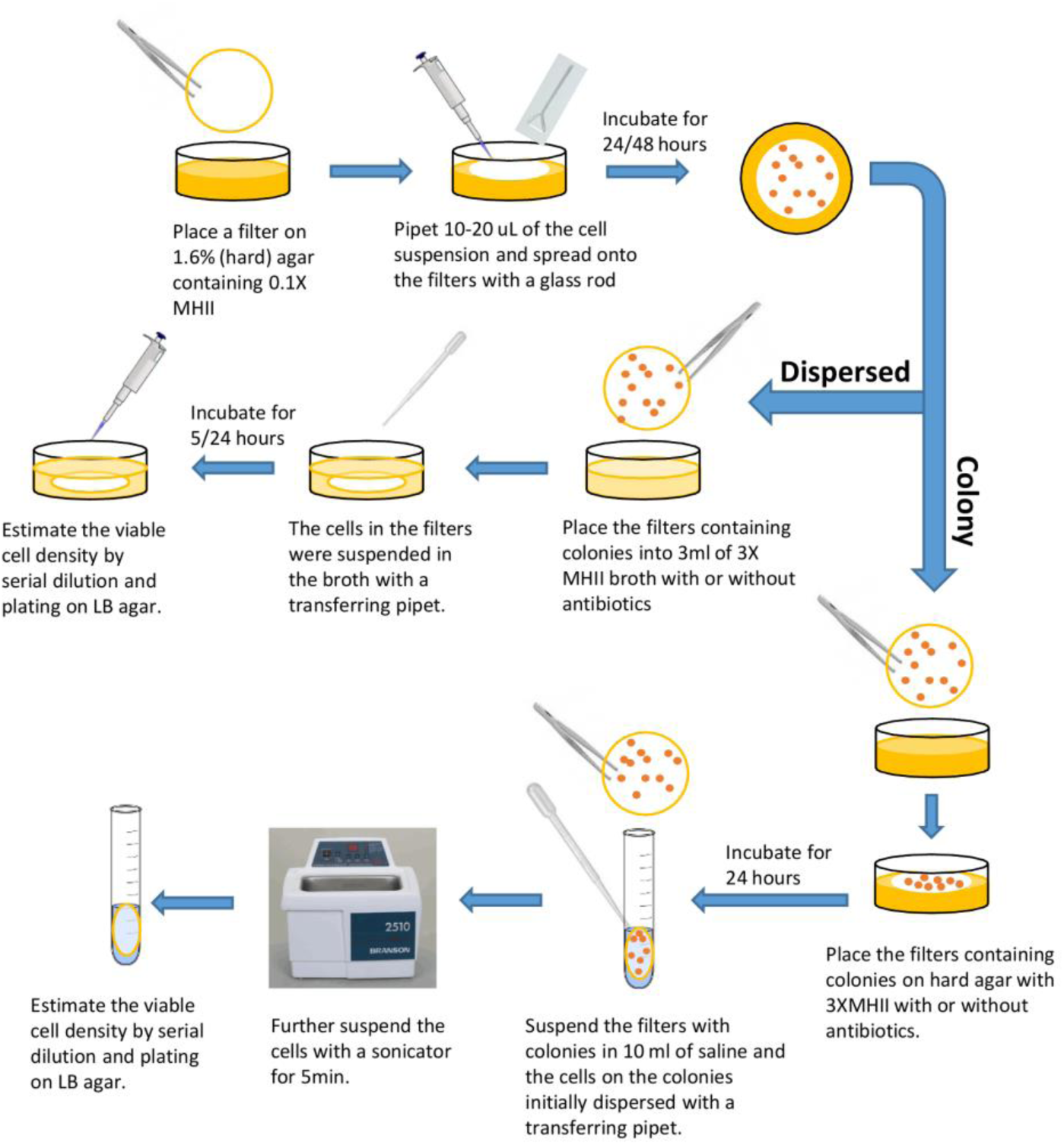
The experiment setup and protocol to grow and treat *S. aureus* Newman grown as colonies on filters. The growth of bacterial colonies is the same for the colony cultures and the dispersed cultures. After 24 or 48 hours of growth, the filters with bacterial colonies are either transferred into fresh liquid broth for the dispersed cultures, or onto fresh agar for the colony cultures.

1. <u>Establishing the colony and liquid cultures:</u> Overnight cultures of *S. aureus* Newman were grown in MHII and serially diluted in 0.85% saline. Using glass rods, the noted numbers of bacteria were spread onto 25mm diameter 0.45 micron Tuffryn^TM^ filters. The filters were placed onto 3 ml of 1.6% agar with 0.1% concentration of standard MHII (2.2 rather than 22 grams per liter) in the wells of Costar Macrotiter 6-well plates. In parallel, the same densities of cells from the overnight culture were put into 9ml 0.1X MHII liquid broth. The filters and the parallel liquid cultures were incubated at 37°C for 24 or 48 hours, the latter with shaking.
2. <u>Exposing the bacteria to antibiotics:</u> (i) For the experiments examining the effects of antibiotics on intact colonies, filters with 24 or 48 hour old colonies grown on 0.1X MHII (2.2 grams per liter) were transferred onto 3 ml of 1.6 agar media containing either standard 1X MHII or 3X MHII (respectively 66 grams per liter) with either 10X or 40X MIC of the antibiotic as well as an antibiotic-free controls. The colony cultures were incubated at 37°C for an additional 24 hours. (ii) For the experiments with planktonic cells growing in liquid the 24 or 48-hour cultures in 0.1X MHII were diluted to the approximate density of bacteria in the corresponding colony cultures and then put into flasks containing either 1X MHII or 3X MHII medium with 10X MIC of the antibiotics or antibiotic-free controls. These liquid cultures were incubated for 24 hours with continuous shaking. (iii) For the bacteria dispersed from the colonies, the filters with 24 or 48 hour old colonies were transferred into 3 ml of 3X MHII broth with 10X MIC ciprofloxacin (DIS-CIP) or oxacillin (DIS-OXY) or without antibiotics (DIS-CON). These cultures were then incubated at 37℃ with continuously shaking for 5 or 24 hours.
3. <u>Sampling and estimating the viable cell densities:</u> The liquid and the dispersed colony cultures were sampled directly by serial dilution and plating on LB agar. The filters with colonies were removed from the agar, placed in 10ml of 0.85% saline and the cells were washed and suspended in saline with plastic Pasteur Pipettes. To suspend the cells and break-up clumps the saline containing filters and cells were sonicated (Bronson, 2510R-DTH, output 100W, 42 kHz±6%) for 5 minutes and vortexed for 10 seconds before serial dilution.

## RESULTS

### The growth of *S. aureus* Newman in liquid and as colonies

The pharmacodynamics of antibiotics and bacteria depend on the physiological state of the cells, which is reflected in their growth rates and the densities of their population (25). To explore these effects and determine the contribution of the physical structure of the population the growth of *S. aureus*, we estimated the densities of cultures growing as colonies with that of cultures of the same initial density growing in liquid (Fig. 3).

**Figure 3.**
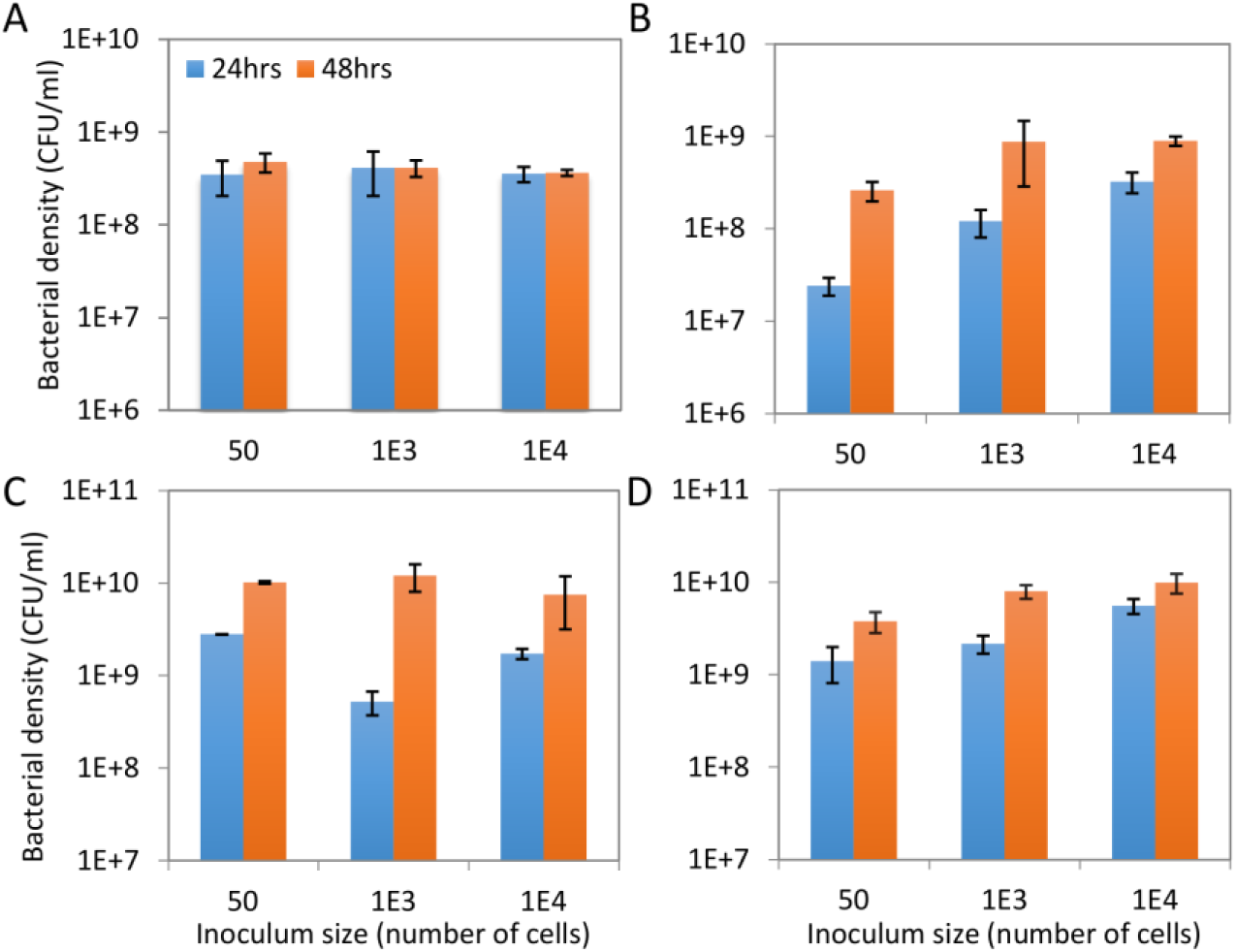
Density and time before resource saturation of liquid and colony populations of *S. aureus* Newman. Cultures were inoculated with 50 cells, 103 cells and 104 cells respectively. Viable cell density of *S. aureus* Newman at 24 (blue bar) and 48 (orange bar) hours grown as (A) Planktonic cells in liquid culture with 0.1X MHII, (B) Colonies on filters on agar with 0.1X MHII, (C) Planktonic cells in 3X MHII broth, and (D) Colonies on filters on 3X MHII agar. Error bars are SEMs.

In liquid culture with 0.1X MHII there is no significant effect of the initial inoculum on the viable cell density of *S. aureus* Newman at 24 hours (P ~ 0.9) (Fig. 2A). On the other hand, with 3X MHII in liquid, at all inoculum densities the bacteria are still growing at 24 hours. At 48 hours in liquid there is no effect of the inoculum density in either 0.1X MHII or 3X MHII (P>0.05) (Fig. 2A and C). As colonies, at all inoculum densities, the population is still growing at 24 hours on both 0.1X MHII and 3X MHII agar (Fig. 2B and D). At 48 hours in both 0.1X MHII and 3X MHII the population inoculated with 50 cells and thereby larger colonies is still growing, whilst those inoculated with 10^3^ and 10^4^ cells appear to be at stationary phase (p>0.05).

**Figure 2.**
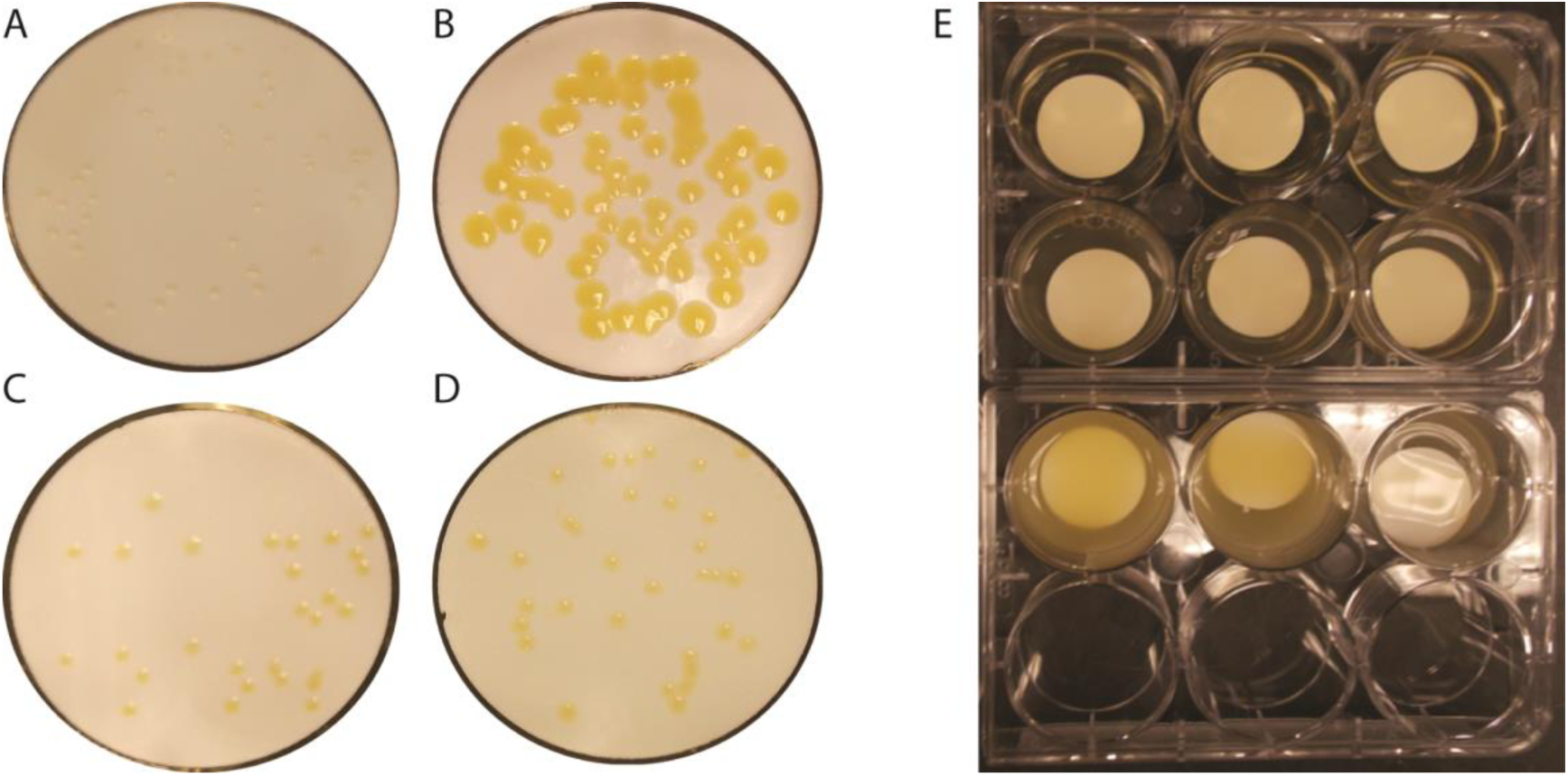
*S. aureus* Newman colonies grown on filters, roughly 50 colonies on each filter. (A) Colonies on filters grown for 48 hours on 3 ml of agar media with 0.1X MHII. (B) 48 hour colonies placed on 3ml agar with 3X MHII for an additional 24 hours. (C) A filter with 48 hour colonies placed on 3ml 3X MHII agar with 10X MIC oxacillin for an additional 24 hours. (D) A filter with 48 hour colonies placed on 3ml 3X MHII agar with 10X MIC ciprofloxacin for an additional 24 hours.

We interpret these results to mean that when liquid cultures in 0.1X MHII are exposed to antibiotics, they are already at stationary phase at 24 hours, but in 3X MHII they are still growing. At 48 hours with inoculates of 10^3^ and 10^4^ cells, these colonies are no longer growing and presumably the cells are at stationary phase, which is not the case for cultures initiated with 50 colonies. It should be noted, that at stationary phase the density of *S. aureus* growing as colonies on 0.1X MHII agar is significantly greater than that in the corresponding liquid culture (p<<0.05 for 10^3^ C and 10^4^ C) (also see (26)) for similar observations). This is not the case for the richer media, 3X MHII, where there is no significant difference in the 48 hour estimated densities when the bacteria are growing in liquid or as colonies (p>>0.05).

### The effect of physical structure on the susceptibility of colonies to antibiotics

We initiate our consideration of the relative antibiotic susceptibility of *S. aureus* growing as colonies and liquid with two bactericidal antibiotics, ciprofloxacin and oxacillin. To determine the contribution of the physical structure of the colonies to these antibiotics, we compare the extent of antibiotic-mediated killing of *S. aureus* in colonies with that of planktonic cells released from colonies (dispersed, DIS) and liquid cultures of the same age as the colonies (LIQ). All were treated with 3X MHII broth or agar containing 10X MIC of these drugs and the viable cell densities were estimated after 5 and 24 hours of exposure to the antibiotics. In Fig. 2, we illustrate the experimental setup and the effects of exposure on the size of 48-hour colonies in the absence of antibiotics and following exposure to oxacillin and ciprofloxacin.

For this assay, we estimated the viable cell density of *S. aureus* at 5 and 24 hours of exposure to oxacillin and ciprofloxacin of cells maintained in colonies (COL), cells released from colonies (DIS) and cells maintained in liquid cultures (LIQ) of the same age (Fig. 4). As noted earlier, both the 24-hour and 48-hour control cultures (CON) continued to increase in density after being transferred to the richer media (Fig. 4) and the size of colonies and the pigmentation dramatically increased (Fig. 2). For ciprofloxacin, 24-hour old planktonic cells grown in liquid culture or released from colonies were more susceptible to killing by this antibiotic after 5 hours of exposure than those maintained in colonies. After 24 hours of exposure, the bacteria within colonies or dispersed from colonies appeared more sensitive to killing by this fluoroquinolone than those in liquid culture (p<< 0.05). After 48 hours of growth, the bacteria within colonies are more refractory to ciprofloxacin than they are as planktonic cells. For both the 24 and 48-hour cultures, relative to the antibiotic-free controls, ciprofloxacin was effective in preventing the growth and killing *S. aureus* with colonies.

**Figure 4.**
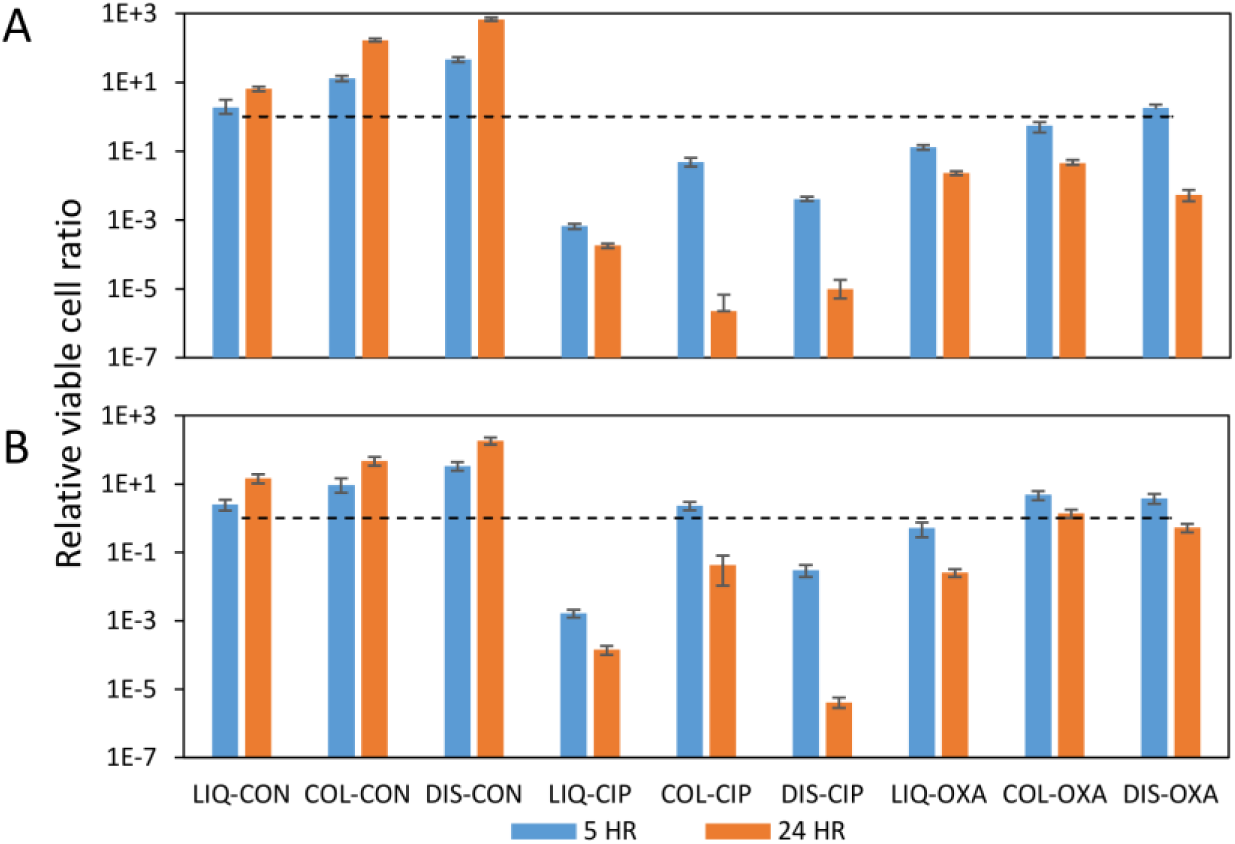
Relative survival of *S. aureus* exposed to 10XMIC ciprofloxacin (CIP) or oxacillin (OXA), in liquid (LIQ), as planktonic cells suspended from colonies (DIS), and as intact colonies (COL). An average of 50 colonies was inoculated on each filter. The CON cultures are antibiotic-free controls. (A) Cultures grown for 24-hour before exposure to the antibiotic. (B) Cultures grown for 48 hours before exposure to the antibiotics. The viable cell densities were estimated at 5 hours and 24 hours of exposure, respectively the blue and red bars. The dashed lines are the viable cell densities before exposure to the antibiotics. Means and standard errors for three replicates.

Oxacillin was clearly less effective than ciprofloxacin in killing *S. aureus* in planktonic cells as well as within colonies. Moreover, there appeared to be little difference in the efficacy of this beta-lactam antibiotic in killing *S. aureus* in colonies relative to that of planktonic cells in liquid. This experiment was repeated three times and similar results obtained (data available upon request).

### Effect of colony size and density on the extent of antibiotic-mediated killing

How does the size (number of cells within and the physical dimensions) of colonies affect their susceptibility to killing by antibiotics? How does the distance between colonies, the density on the agar affect their susceptibility to killing by antibiotics? Most importantly, how effective are different antibiotics in inhibiting the growth and killing of *S. aureus* as planktonic cell liquid and within colonies. To address these questions, we prepared filters with ~ 50 cells and ~ 10^4^ cells, respectively large, widely dispersed colonies and small, densely distributed colonies. These filters were placed on 0.1X MHII and grown for 24 or 48 hours, at which time they were placed on 1X MHII agar containing 10X MIC of one of six bactericidal antibiotics or one of three bacteriostatic drugs.

In Table 2 we present the results of these experiments with cultures initiated with approximately 50 colonies. In the absence of antibiotics in both liquid and as colonies, the bacterial population increased in density. The bacteriostatic drugs, tetracycline erythromycin and linezolid prevented the population from growing in colonies as well as in liquid. The most effective bactericidal antibiotic in killing both planktonic *S. aureus* and with within colonies was ciprofloxacin. For the younger, 24 hour cultures this fluoroquinolone was more effective in killing the bacteria in colonies than in liquid, the opposite was true for the older 48 hour cultures. For the younger cultures gentamicin was similarly effective in killing the cells in colonies as it was when these bacteria were in liquid. This is not the case when the cultures were 48 hours old at the time of exposure to this drug, which was ineffective in killing these bacteria when they were in colonies.

**Table 2.**
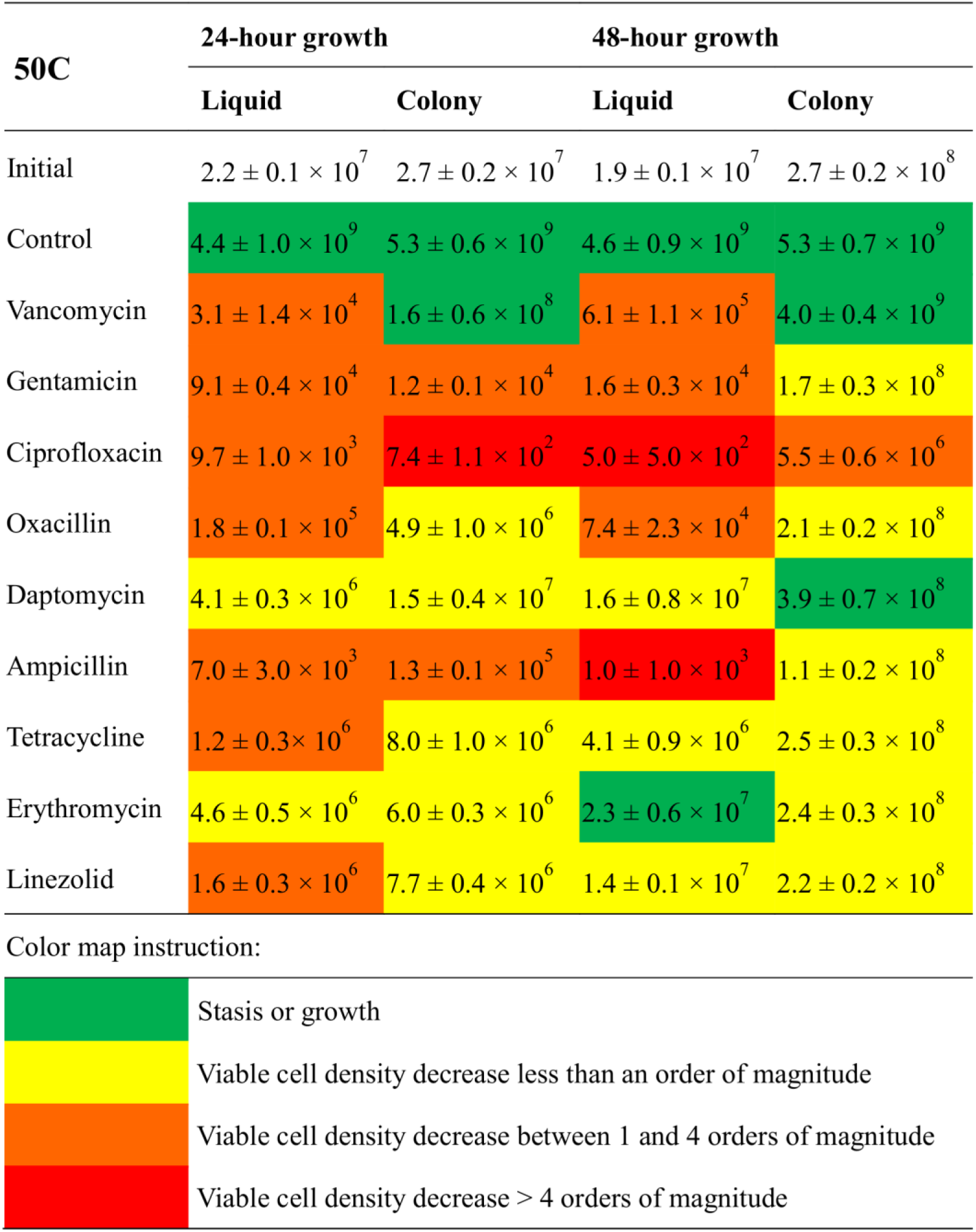
The results of experiment with *S. aureus* Newman cultures initiated with an average of 50 cells per 3ml of MHII broth/agar containing 10X MIC of the drugs listed. Cultures grew for 24 or 48 hours in 3ml of 0.1X MHII broth/agar before exposure to the antibiotics in 1X MHII. The viable cell densities (CFU/ml) were estimated 24 hours after exposure to the antibiotics.

Vancomycin was moderately bactericidal in liquid but totally ineffective when bacteria were in colonies. At these densities, daptomycin was effectively bacteriostatic in both liquid and younger colonies, but failed to prevent the growth of *S. aureus* the 48hour old colonies. This may be well attributed to the relatively high initial density of the culture and the degradation of this drug at this higher cell densities (27, 28). The beta-lactam antibiotics were less effective in killing *S. aureus* in colonies than they were when these cells were in liquid.

In the cultures inoculated with ~10^4^ cells the colonies were much smaller and more crowded than the colonies inoculated ~ 50 cells per filter (Table 3). Of the 9 antibiotics considered, gentamicin and ciprofloxacin were the most effective at killing *S. aureus* when treating the 24-hour cultures, even more effective than they were with the larger colonies. Apart from gentamicin and ciprofloxacin, all the other antibiotics were less effective comparing to corresponding cultures inoculated ~ 50 cells per filter. The bacteriostatic drugs inhibited the growth of both 24 and 48 hour colonies but failed to do so in all liquid cultures. Vancomycin failed to kill the bacteria in all cases. In general, the 48-hour crowded colony cultures were the most refractory to antibiotics.

**Table 3.**
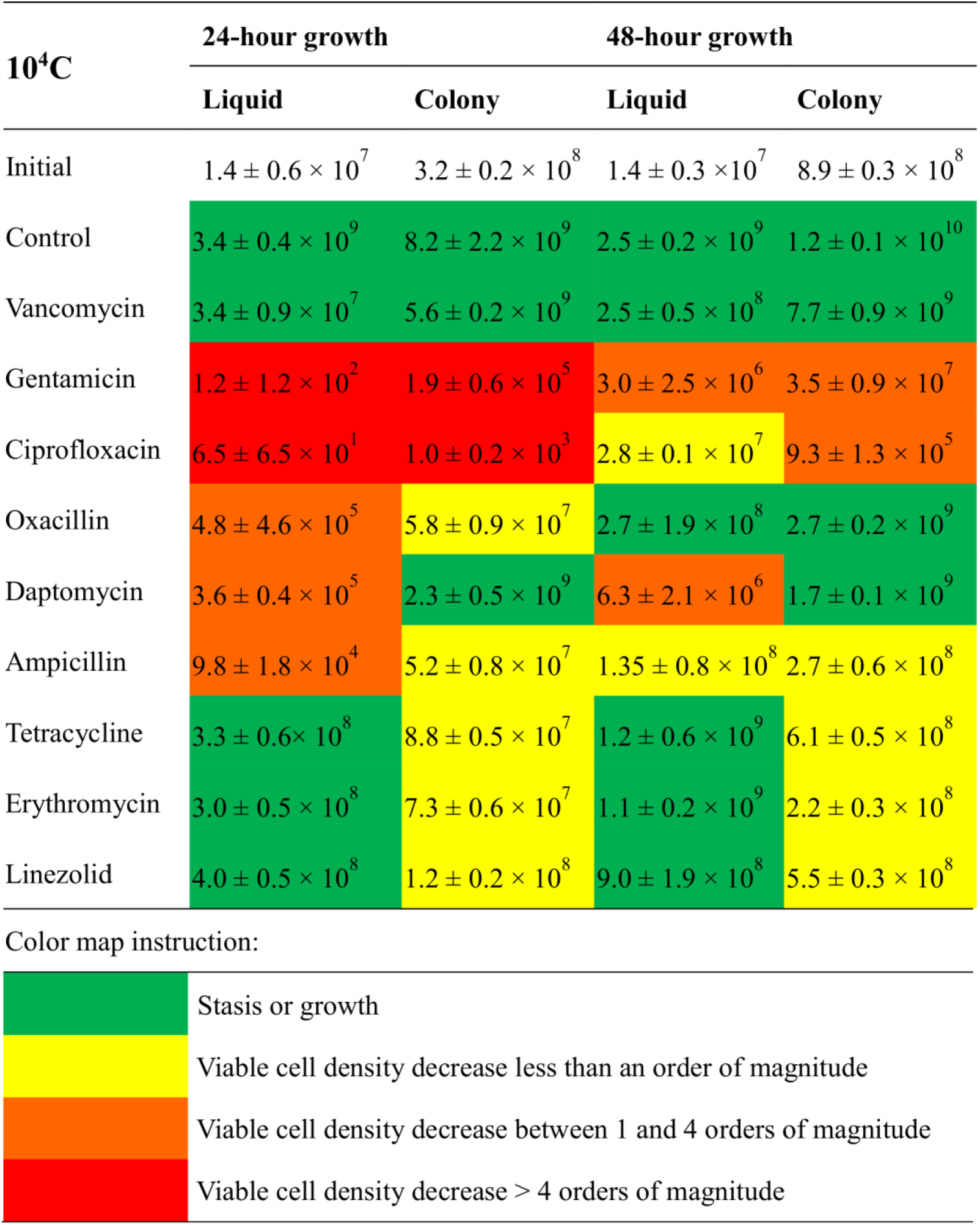
Antibiotic-mediated killing of *S. aureus* Newman cultures initiated with an average of 1e4 cells per 3ml of MHII broth/agar containing 10X MIC of the drugs listed. Cultures grew for 24 or 48 hours in 3ml of 0.1X MHII broth/agar before exposure to the antibiotics. Viable cell density (CFU/ml) was measured at 24 hours after exposure.

## Discussion

In designing and performing these experiments we had two goals. One, to develop and evaluate a facile and broadly applicable method for quantitative studies of the antibiotic susceptibility of bacteria growing as colonies. Two, to apply this procedure to explore the efficacy of different antibiotics to inhibit the growth and kill *Staphylococcus aureus* in colonies. While there have been a number of studies of the efficacy of antibiotics for treating bacteria in biofilms (17–22, 29–32), to our knowledge this is the first investigation to explore antibiotic-mediated inhibition of replication and killing of bacteria growing as discrete colonies on surfaces.

The procedure developed here can be applied to virtually any bacteria that when grown in vitro forms colonies. This same procedure could be employed to evaluate the susceptibility of single and multi-species biofilms to antibiotics. For this, instead of seeding the filters with relatively few bacteria to form discrete colonies, the filters could be seeded with large numbers of bacteria of the same or multiple species. The sampling methods would be identical to those described in here. By comparing liquid cultures and bacteria released from colonies (or biofilms) of the same density and stage of growth (age) this method provides a way to evaluate the contribution of the physical structure of the population to its susceptibility to antibiotics.

The results of this study indicate that there is substantial variation among bactericidal antibiotics in their efficacy for killing bacteria within colonies. Of the six bactericidal antibiotics considered here, ciprofloxacin was most effective, followed by gentamicin. Of the beta-lactam antibiotics, ampicillin was more effective in killing *S. aureus* in colonies than oxacillin. In our experiments, daptomycin, which is considered bactericidal (33–35), was no more capable of killing *S. aureus* in colonies than the antibiotics that are deemed bacteriostatic, tetracycline, erythromycin and linezolid. Indeed, this cyclic peptide was less effective in preventing the proliferation of *S. aureus* in more mature (48 hour) colonies than these bacteriostatic drugs. Vancomycin, which is commonly employed for treating methicillin resistant *S. aureus* (7, 9, 32, 34, 36), was virtually ineffective for either preventing the replication of or killing *S. aureus* Newman in colonies.

The preceding conclusions about the relative efficacy of the different antibiotics for treating *S. aureus* as colonies is based on a common dose of 10X MIC of these drugs with 24 and 48-hour inoculum densities respectively of ~2×10^7^ and ~2×10^8^ cells per ml for the experiments with 50 colonies and ~3×10^8^ and ~9×10^8^ for the experiments initiated with 10^4^ colonies. These densities are substantially greater than the recommended 5x105 cells per ml for estimating MICs (24) by serial dilution. To some extent the differences in relative efficacy of the bactericidal antibiotics to kill *S. aureus* in colonies may be attributed to a density (inoculum) effect (27). On the other hand, we don’t see the utility of reducing the density of cells treated to make these drugs more effective in this experimental system. From a clinical perspective, concern is to treat established infections the densities of which are likely greater than the 5×10^5^ (37, 38).

What about increasing the dose of the antibiotics that were ineffective for treating colonies in these experiments? We have explored this possibility with 40X MIC of oxacillin and vancomycin. The results of these experiments suggest that even at these high concentrations, these antibiotics are ineffective for treating *S. aureus* Newman in colonies (Table S1).

On first consideration it may seem that the relative inefficacy of vancomycin, oxacillin and daptomycin to kill or prevent the replication of *S. aureus* in colonies may reflect the inability of these drugs to diffuse through the membrane on which the colonies are growing and then through the colonies. We do not believe this is the case for oxacillin or vancomycin. Although it is known that to some extent biofilms reduce the rate of diffusion of antibiotics. In our experiments, however, the time of exposure to these drugs was relatively long, 24 hours. We expect that the diffusion effect would be small. Based on what is known about the diffusion rates of these drugs in *Staphylococcus* in biofilms it seems reasonable to assume the bacteria within these colonies would have been exposed to substantial concentrations of these drugs (30, 31, 39). Moreover, the cells dispersed from colonies before treatment with these drugs were no more susceptible than those in intact colonies, even though the bacteria were in liquid and confronted with the same concentration of these antibiotics (Table S1). We suggest the primary reason for the relative inability of these antibiotics to kill *S. aureus* in colonies can be attributed to the density and physiological state of the bacteria, rather than the structure of the colonies.

One possible explanation for why daptomycin is not effective in colonies but is in liquid is that its mode of action is thwarted by the structure of the colonies. It has been proposed that daptomycin operates by depolarizing the cell membrane which results in leakage of ions (7–9, 30). If, however, the cells are within colonies the ions lost by individual cells would remain in the collective and thereby shared by all in this community.

## Acknowledgements

This work was supported by a grant from the US National Institutes of Health, GM 091875 (BRL) and Laney Graduate School of Emory University.

